# Test-retest reliability of tone- and 40 Hz train-evoked gamma oscillations in female rats and their sensitivity to low-dose NMDA channel blockade

**DOI:** 10.1101/2021.01.08.425905

**Authors:** Muhammad Ummear Raza, Digavalli V. Sivarao

## Abstract

**Rationale:** Schizophrenia patients consistently show deficits in sensory-evoked broadband gamma oscillations and click-evoked entrainment at 40 Hz, called the 40 Hz auditory steady-state response (ASSR). Since such evoked oscillations depend on cortical N-methyl D-aspartic acid (NMDA)-mediated network activity, they can serve as pharmacodynamic biomarkers in the preclinical development of drug candidates engaging these circuits. However, there is little test-retest reliability data in preclinical species, a prerequisite for within-subject testing paradigms.

**Objectives:** We investigated the long-term stability of these measures in a rodent model.

**Methods:** Female rats with chronic epidural implants were used to record tone- and 40 Hz click-evoked responses at multiple time points and across six sessions, spread over 3 weeks. We assessed reliability using intraclass correlation coefficients (ICC). Separately, we used mixed-effects ANOVA to examine time and session effects. Individual subject variability was determined using the coefficient of variation (CV). Lastly, to illustrate the importance of long-term measure stability for within-subject testing design, we used low to moderate doses of an NMDA antagonist MK801 (0.025-0.15 mg/kg) to disrupt the evoked responses.

**Results:** We found that 40 Hz ASSR showed good reliability (ICC=0.60-0.75) while the reliability of tone-evoked gamma ranged from fair to good (0.33-0.67). We noted time but no session effects. Subjects showed a lower variance for ASSR over tone-evoked gamma. Both measures were dose-dependently attenuated by NMDA antagonism.

**Conclusion:** Overall, while both measures use NMDA transmission, 40 Hz ASSR showed superior psychometric properties of higher ICC and lower CV, relative to tone-evoked gamma.

## Introduction

Schizophrenia is a disorder in which information processing is abnormal at multiple levels, from distorted transmission to impaired perception (Javitt, 2009a, 2009b; Krishnan et al., 2011; Silverstein and Keane, 2011). Sensory-evoked neural responses in patients tend to be smaller and less synchronized across many paradigms (Javitt, 2009a; Luck et al., 2011; Rissling and Light, 2010), suggesting an abnormal registration and/or processing. Specifically, discrete auditory stimulus-evoked oscillations in the gamma frequency range (30-100 Hz) are consistently abnormal in schizophrenia as well as in other neuropsychiatric conditions like autism and bipolar disorder. The local interactions between parvalbumin-positive GABAergic basket cells and pyramidal cells are important for the generation of gamma oscillations (Gonzalez-Burgos and Lewis, 2008; Tiesinga and Sejnowski, 2009). Specifically, pyramidal neuron feed-forward activation of the N-methyl D-aspartic acid (NMDA) receptors on the basket cells and their feedback inhibition of the pyramidal cells is critical for the emergence of these oscillations (Buzsáki and Wang, 2012; Cardin et al., 2009; Gonzalez-Burgos & Lewis, 2012).

Principally, two types of gamma oscillations are evoked by auditory stimuli like tones and clicks. A transient burst of broad-band gamma oscillatory activity (~ 30 Hz to 100 Hz) that is coincident with the auditory evoked P1-N1-P2 potential (Javitt and Sweet, 2015; Ross et al., 2010) is observed in response to a discrete tone or a click stimulus. A second type of response, called the auditory steady-state response (ASSR), is an electroencephalographic (EEG) entrainment reflecting the driving frequency of a periodic auditory stimuli such as clicks or amplitude-modulated tones, lasting as long as the train stimulus, and typically elicited at 40 Hz (Brenner et al., 2009). Both these responses are deficient in not only schizophrenia patients but also individuals that are at high risk for the disease (Leicht et al., 2011, 2016; Oribe et al., 2019; Perez et al., 2013), making them state and trait markers. Moreover, such sensory gamma deficits correlate with symptoms as well as in some studies, long-term functional outcomes for the patients (Molina et al., 2020). Thus, gamma oscillatory measures may reflect abnormalities in key cortical circuits involved in symptoms and impacting clinical outcomes. There is now a growing interest in using evoked gamma oscillations as a functional biomarker for neuropsychiatric drug development (Kozono et al., 2019; Light et al., 2020; Luck et al., 2011; Sivarao, 2015; Sivarao et al., 2016).

In psychopharmacological drug development, there is now a greater appreciation for patient heterogeneity and its contribution to high odds ratios seen with many clinically efficacious drugs (Stroup et al., 2007). Clinical trials that use “within-subject” designs tend to better control such heterogeneity over “between-subject” designs (Salkind, 2010). More recently, within-subject designs involving the so-called N=1 trials are emerging as a sensitive means for identifying the most effective treatment for an individual patient, taking into account his or her unique genetic and epigenetic backgrounds (Lauschke et al., 2019). However, stable, objective circuit-level biomarkers can de-risk or accelerate such efforts, by enabling proper dose selection and demonstrating neurophysiological engagement.

In preclinical discovery too, biomarkers are critical to demonstrate circuit engagement and enable prioritization of molecular leads for further development. Here too, within-subject crossover designs are more sensitive to pharmacological treatment, yielding typically better effect sizes over between-subject designs, and requiring smaller group sizes to demonstrate treatment effect (Cleophas and De Vogel, 1998; Guo et al., 2013).

However, the use of such biomarkers in within-subject design studies is predicated on their temporal stability, to support multiple levels of drug treatment lasting weeks, if not months. Several recent studies have reported the stability of 40 Hz ASSR measures in healthy human subjects (McFadden et al., 2014) as well as in schizophrenia patients (Cervenka et al., 2013; Ip et al., 2018; Legget et al., 2017; McFadden et al., 2014; Roach et al., 2019; Roach et al., 2019). However, to the best of our knowledge, similar studies in preclinical species are not available.

In the current report, we tested the stability of tone-evoked broad-band gamma (30-100 Hz) and 40 Hz click train-evoked narrow-band gamma (35-45 Hz) activity in a group of female rats, in six sessions, spread over a 3-week period. We investigated the test-retest reliability using the intra-class correlation coefficients (ICC) at discrete time points across sessions. To rule out carryover effects of repeated testing, we used mixed-effects ANOVA, using test sessions and time of testing as fixed effects and subjects as a random effect. To estimate the variance at the level of individual subjects, we calculated coefficients of variation (CV). Lastly, to illustrate the importance of stability to pharmacological testing, we used a repeated measures design to test low to moderate doses of the high affinity, open NMDA channel blocker MK-801, to disrupt evoked oscillations.

## Methods

### General

The Institutional Animal Care and use Committee of the East Tennessee State University approved all experimental procedures involving animal subjects. Female Sprague Dawley rats (6-8-week-old; 200-250 g) were obtained from Envigo (Indianapolis, IN) and group housed until surgery, with free access to food and water.

### Surgical

Surgeries were performed under isoflurane anesthesia (5% induction; ~ 2% maintenance, in oxygen; 1 liter/min flow rate). The head was secured in a stereotaxic frame and core temperature was maintained at 36±1^0^ C. Head was shaved and disinfected with alcohol and povidone-iodine swabs. Bupivacaine (0.25 %, sc) was used for a local nerve block and ketoprofen (5mg/kg, sc) was used for perioperative analgesia. Atropine sulfate (0.05 mg/kg, sc) was given to check respiratory secretions. The skull was exposed through a midline incision, followed by blunt dissection. A solution of 3% hydrogen peroxide solution was used to disinfect and dry the skull. An aseptic drill bit was used to drill four burr holes into the skull and epidural electrodes with attached wires were screwed in, avoiding damage to underlying dura. Electrode coordinates were as follows: frontal, 1 mm anterior to bregma and 1 mm lateral to midline; vertex, 4.5 mm caudal to bregma and 1 mm lateral to midline; reference, 2 mm caudal to lambda and 2 mm lateral (left side); ground, 2 mm caudal to lambda and 2 mm lateral (right side). Dental cement slurry was poured over the electrodes and allowed to set. Electrode wires were soldered to the recording head mount (Pinnacle Technology, Lawrence, KS). For EMG, two wires from the head mount were sutured to the nuchal muscle. The area was covered with a second layer of dental cement and after curing, was closed using silk sutures. Rats were allowed to recover from surgery for at least 10 days and were later handled and acclimated to the recording set up before use in any experiments.

### EEG recording

Rats were placed in Plexiglas cylinders (Pinnacle Technology, Lawrence, KS) equipped with a video camera and a house speaker (DROK, B00LSEVA8I). Rats were tethered to the recording setup through a shielded preamplifier cable and a commutator for continuous EEG recording, while permitting free access to explore within the chamber. The acquisition system (CED Power 1401; Cambridge Electronics Design (CED), Cambridge, UK) was used to produce auditory stimuli, as well as to acquire the EEG (Signal v7). Data were acquired as 5 s sample sweeps (1 KHz). A tone stimulus was presented at 1 s after a sweep initiation (1 KHz, 2mV sinusoidal wave, 50 ms duration, 65 dB). A click train (5 mV monophasic 1 ms long square waves, 20/0.5 s, 65 dB) was presented 2 seconds after the tone stimulus. Interval between the end of the click train and the beginning of the next tone was 2.5 s. For the ICC study, data were acquired at 0, 30, 60, 90- and 120-minutes. The 0 min time point was omitted in the pharmacology study. Six sessions, each separated by at least 3 days, was used for the reliability study, whereas pharmacology study was carried out in four sessions.

### EEG data analysis

While all EEG data were evaluated for movement artifacts, generally there were very few frames (<3%) that needed exclusion. Artifact free frames were averaged for each subject. Averaged EEG was filtered (30-100 Hz for tone and 35-45 Hz for 40 Hz click-train using a second order Hanning infinite impulse response (IIR) filter; Signal v7). Root mean square (RMS) amplitudes were computed for tone-evoked (0-150 ms from tone onset) and 40 Hz ASSR (0-500 ms, from train onset) responses, for each subject, at each time point. These were further averaged to summarize group responses.

### Pharmacology

MK801 maleate (MW 337.4 g/mol) was obtained from Sigma-Aldrich. Free base mass was used to make 0.15 mg/kg stock solution and this was diluted to make 0.025 and 0.05 mg/kg. These doses were chosen to capture a wide range of NMDA channel occupancy (Fernandes et al., 2015). Rats were given intraperitoneal (IP) injection of either vehicle or MK-801 at the start of experiment and were then placed in the recording chamber. At least 3-day period was allowed for washout, between treatments. Animals were carefully monitored for any carryover effects and weight fluctuations.

### Statistics

Since a key goal of this work was to evaluate the suitability of evoked gamma oscillations as biomarkers in repeated measures design, we wanted to know if there were any systemic effects of repeated testing. For this, we used a mixed effects ANOVA (restricted maximum likelihood estimation as implemented in Graphpad Prism v8.4.3 software). Time of testing within a session was designated as a row factor, while multiple sessions were designated as a column factor. Data normality was assessed using quantile-quantile (q-q) plots. Sphericity was not assumed and Greenhouse-Geisser correction was used. Where appropriate, Dunnett’s tests were used to evaluate within session significance, using time zero response as a comparator. For pharmacology studies, vehicle group was used as a comparator.

### Reliability was measured by calculating intraclass correlation coefficients (ICC)

There are many types of ICC reported in the literature. We implemented the two-way mixed effects, absolute agreement, single measurement formula, as discussed by Koo and Li (2016). ICC estimates along with their 95% confidence intervals were calculated using SPSS statistical package (SPSS v26, Chicago, IL). We used a qualitative interpretation of the ICC values as reported recently by Ip et al., (2018). Thus, an ICC less than 0.39 was considered to be an indicator of poor test-retest reliability, while ICC ranges between 0.40-0.59, 0.60-0.75 and 0.76 and above, were considered respectively to be fair, good or excellent (Ip et al., 2018).

## Results

Auditory stimuli of tone and click train, evoked broadband gamma and steady-state entrainment responses respectively, in all subjects. An overlay in Figure 1 shows the grand averaged RMS amplitude data from six sessions, corresponding to discrete time points (0 to 120 min). Panel A shows tone-evoked gamma band activity and panel B shows the ASSR response.

**Figure 1.**
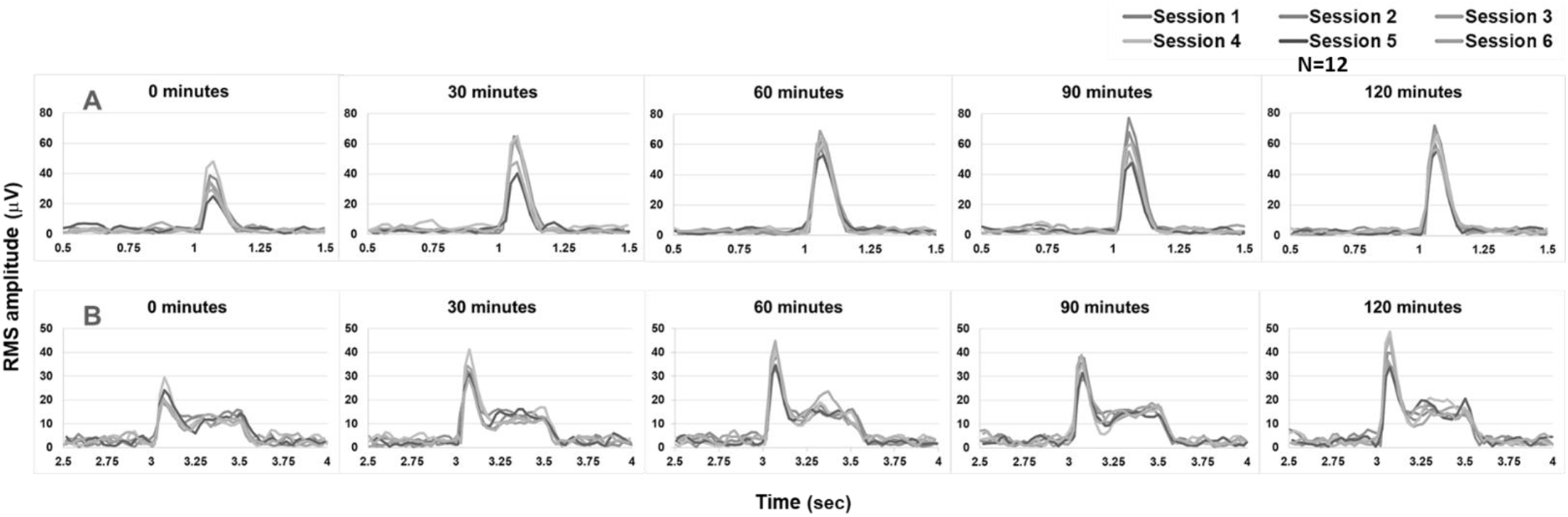
Overlay of the RMS amplitude measure of 1 KHz tone-evoked broad-band gamma (top panel) and 40 Hz click-evoked narrow-band steady-state response (bottom panel) at 0 min −120 min, from 6 discrete sessions, spread over a 3-week period. Data were from a group of 12 rats.

For tone-evoked γ, a linear Q-Q plot confirmed normal distribution (data not shown). Mixed effects ANOVA showed a highly significant effect of time on group means (p<0.0001; F (2.580, 28.38) = 13.91; epsilon = 0.6449). However, no significant session effect (p=0.2195; F (2.458, 27.04) = 1.590; epsilon = 0.4917) or interaction of time x session (p=0.1896; F (5.319, 55.85) = 1.538; epsilon = 0.2659) were noted. Examining the mean response across time indicated a pattern of smaller average responses at time zero relative to other times points (Figure 1, panel A). Confirming this, post hoc analysis showed that the time zero response was smaller than several other time points within each session (p<0.05; Dunnett’s tests). Such differences were noted in session 1 (p<0.05; 0 min vs. 30, 60 and 90 min), session 2 (0 min vs. 30, 60, 90 and 120 min), session 3 (0 min vs. 30, 60 and 120 min) and session 5 (0 min vs. 120 min).

The 40 Hz ASSR data too were normally distributed, based on a linear Q-Q plot (not shown). As with the tone-evoked gamma, mixed-effects analysis of 40 Hz ASSR showed a highly significant effect of time (p=0.0001; F (3.1, 34.10) = 8.956; epsilon = 0.7751) but no session effect (p=0.3021; F (2.5, 27.5) = 1.268; epsilon = 0.5) or interaction of time x session (p=0.6067; F (3.916, 41.12) = 0.6802; epsilon = 0.1958). As with transient gamma, mean ASSR at time zero tended to be smaller than responses at other time points (Figure 1, panel B). Within session Dunnett’s tests revealed following contrasts to be significant (p<0.05): session 2 (0 min vs. 120 min), session 3 (0 min vs 60 min), session 4 (0 min vs. 60 min and 120 min), session 5 (0 min vs. 60 and 120 min) and session 6 (0 min vs. 60 and 120 min).

We next calculated the ICCs for each time point across sessions using a two-way mixed effects model using single measurement, absolute agreement criteria as discussed elsewhere (Koo and Li, 2016). The ICCs along with their 95% CIs, at designated time points and across 6 sessions, are shown in table 1. Tone-evoked gamma, with the exception of one time point, showed an ICC that had a fair measure of reliability (0.33-0.67). On the other hand, ICC for 40 Hz ASSR showed good (0.60-0.75) reliability. Although we did not do a pair-wise statistical comparison between the two, it is apparent from the data summarized in Table 1 that, for the same group of subjects, 40 Hz ASSR had higher ICCs compared to tone-evoked gamma, in each instance.

**Table 1.** Intra-class correlation coefficients (ICC) across time points from six sessions were calculated using a two-way mixed effects model, single measurement and absolute agreement criteria ((Koo and Li, 2016).

| | Tone-evoked $\gamma$ | | 40 Hz ASSR | |
| --- | --- | --- | --- | --- |
| Time | ICC | 95% CI | ICC | 95% CI |
| 0 min | 0.50 | 0.22-0.83 | 0.75 | 0.51-0.93 |
| 30 min | 0.33 | 0.09-0.72 | 0.66 | 0.39-0.90 |
| 60 min | 0.63 | 0.37-0.86 | 0.70 | 0.47-0.89 |
| 90 min | 0.50 | 0.27-0.77 | 0.60 | 0.37-0.83 |
| 120 min | 0.67 | 0.45-87 | 0.74 | 0.54-0.90 |

Since the same group of 12 rats were repeatedly tested ~ 30 times, spread over 3 weeks, we had an opportunity to calculate % coefficient of variation (CV) as a measure of individual subject’s response precision. Mean CV for tone evoked gamma was 31.6%. In contrast, the mean CV for 40 Hz ASSR was markedly lower at 20.9. Figure 2 summarizes these data as a violin plot, highlighting the marked difference between the two measures (p=0.0001; paired t-test).

**Figure 2.**
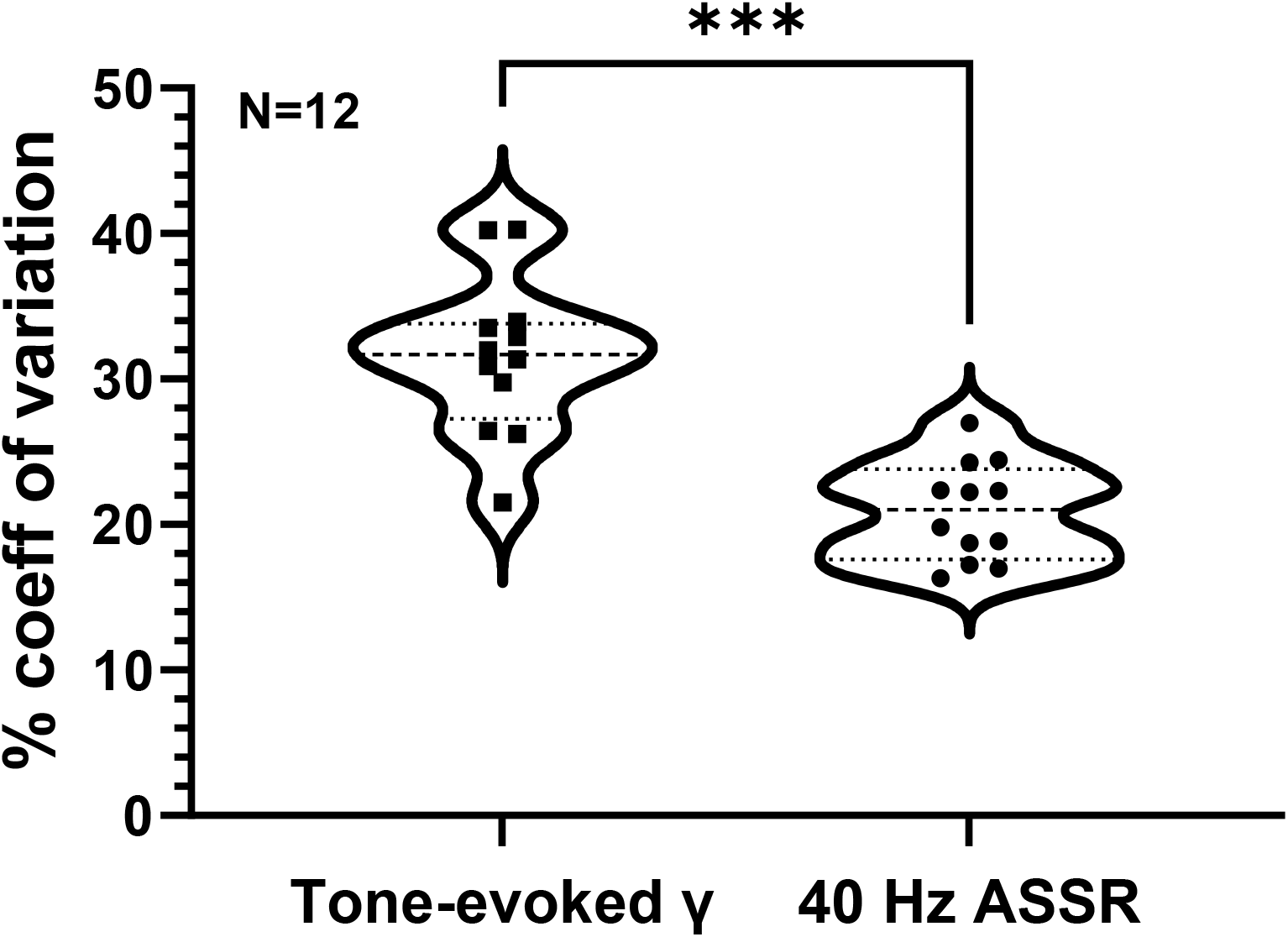
Summary of coefficient of variation (CV) of tone-evoked and 40 Hz click train-evoked responses from a group of 12 rats. Individual subject CVs are indicated within the violin plots. Significant difference between the two groups is indicated by asterisks (P = 0.0001; paired t-test).

In the second part of the study, we illustrated the utility of temporal stability of tone-evoked and train-evoked gamma oscillations, by carrying out a dose-response study using the NMDA channel blocker MK801, using a repeated measures design. We chose to omit testing at time zero, but tested at other time points (30 min −120 min) post-dose. The dose of 0.15 mg/kg dose of MK-801 induced hyperactivity, circumambulation along with stereotypic head movements that persisted for 3-4 hours post-injection. However, no behavioral changes were noted at 0.025 or 0.05 mg/kg doses.

MK801 disrupted tone-evoked gamma in a highly dose-dependent and time dependent manner. Figure 3 summarizes the grand averaged evoked responses for all treatments, across all subjects, 60 min post-treatment, while also showing the individual responses. Figure 4 (top panel) summarizes tone-evoked gamma after vehicle or MK801 treatments. Normal distribution of the data was first ascertained through a Q-Q plot (not shown). Mixed-effects ANOVA showed a robust treatment effect (p<0.0001; F (1.574, 14.16) = 25.98; epsilon = 0.5246) but no time effect (p= 0.4709; F (2.280, 20.25) = 0.8149; epsilon = 0.7600) or time x treatment interaction (p=0.2007; F (2.245, 20.21) = 1.728; epsilon = 0.2495). Post-hoc comparisons using Dunnett’s test showed that only 0.05 and 0.15 mg/kg doses of MK801 significantly (p<0.05) attenuated tone-evoked gamma at multiple time-points, relative to vehicle treatment.

**Figure 3.**
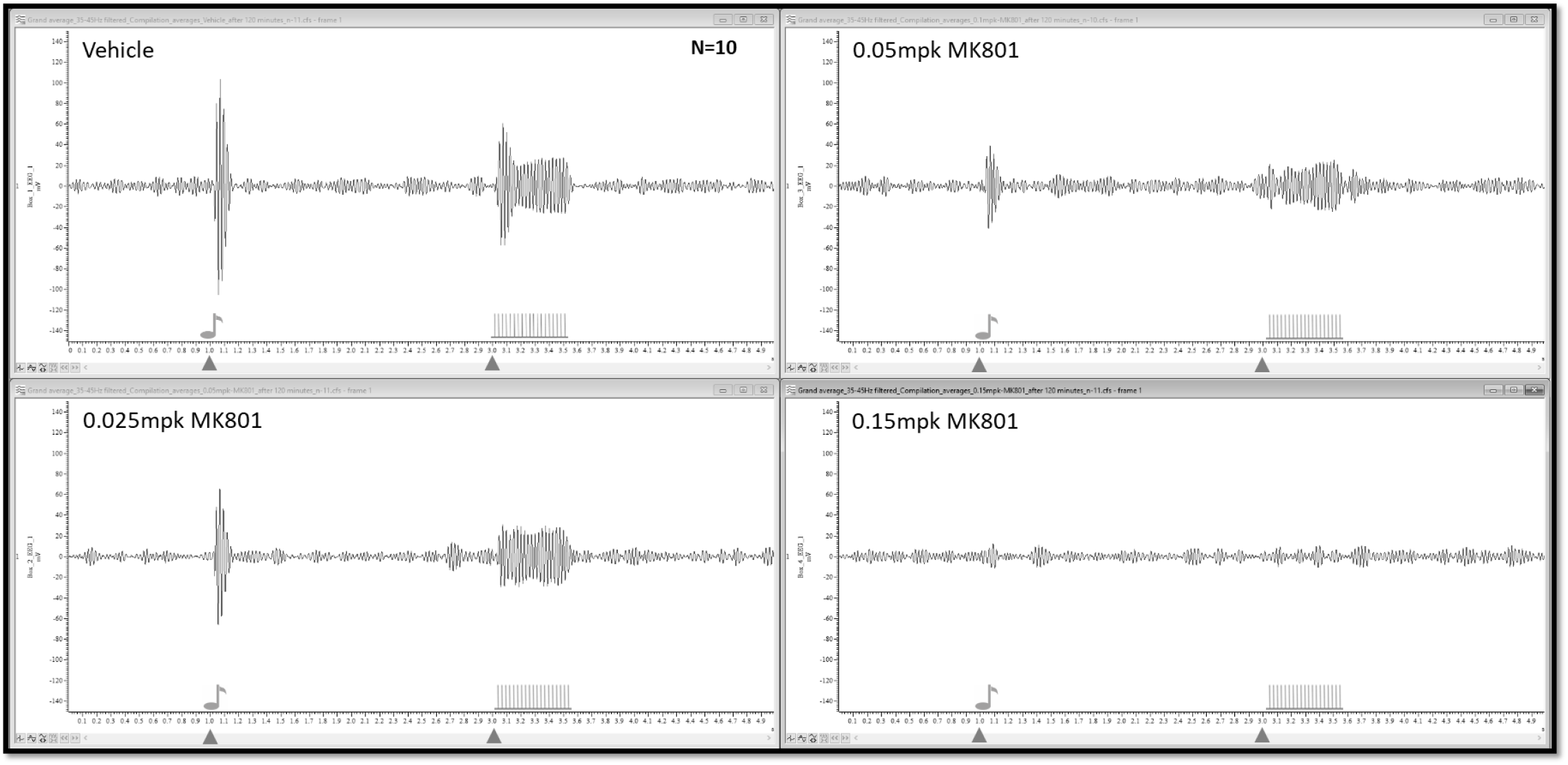
A compilation of grand average of narrow band gamma (35-45 Hz) activity in response to auditory tone (1 KHz) and 40 Hz click stimuli, 120 min after vehicle or MK801 (0.025, 0.05 0r 0.15 mg/kg, ip) treatment. Data represent a grand average from N=10 subjects.

**Figure 4.**
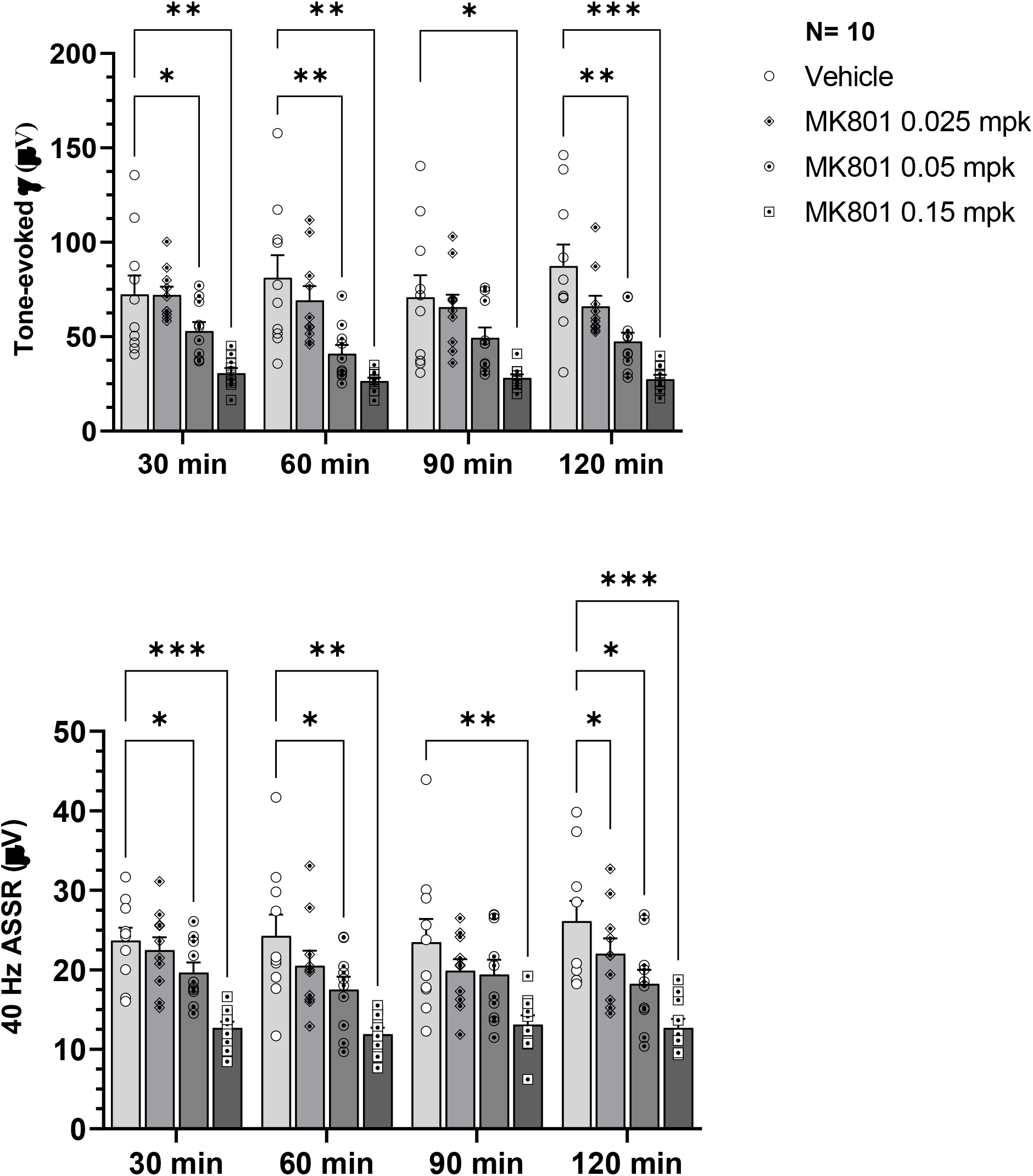
Compilation of RMS data of tone-evoked gamma activity (top panel) or 40 Hz ASSR (bottom panel) in a group of rats treated with vehicle or MK801 (0.025, 0.05 or 0.15 mg/kg sc), at discrete time points (30, 60, 90 and 120 min). Significant differences between vehicle treatment and MK801 treatment are indicated by asterisks (*/**/*** indicates p<0.05/0.01/0.001; Dunnett’s posthoc tests). Data are from a group N=10 rats.

ASSR data too were normally distributed, based on a Q-Q plot (not shown). The 40 Hz ASSR responses are summarized in Figure 4, bottom panel. MK801 dose- and time-dependently disrupted 40 Hz train-evoked gamma. Robust treatment effect was noted (p<0.0001; F (1.884, 16.95) = 31.39; epsilon = 0.6279). However, neither time (p=0.4365; F (2.679, 24.11) = 0.9208; epsilon = 0.8930) nor time x treatment (p=0.5328; F (3.800, 34.20) = 0.7922; epsilon = 0.4222) were significant. Post-hoc testing showed that all three doses were effective (p<0.05) at one or more time points, in reducing 40 Hz ASSR response, compared to vehicle treatment.

## Discussion

Circuit-based functional biomarkers are critical for early phase drug development; for prioritizing lead candidates in preclinical species and in the clinic for dose selection, proof of concept, and patient stratification (Light et al., 2020; Spellman and Gordon, 2015). While there are several recent clinical reports that show test-retest reliability for tone- and train-evoked gamma oscillations (Cervenka et al., 2013; Legget et al., 2017; McFadden et al., 2014; Tan et al., 2015), such findings only infrequently extend beyond a couple of test sessions (Ip et al., 2018; Roach et al., 2019). Moreover, to date, complementary reliability studies of evoked gamma in preclinical species are unavailable. Thus, our current report addresses an important translational knowledge gap by establishing test-retest reliability of tone- and 40 Hz-train-evoked gamma oscillations across sessions, spread over a three-week period.

Since our interest is in using these EEG measures for pharmacological testing using repeated measures design, we evaluated time and session effects (fixed variables) with subjects modeled as a random effect, using a mixed effects analysis. We found a robust time effect with time zero responses tending to be smaller in comparison to all other time points. Although rats were extensively acclimated, transfer from home cage to the recording chamber triggered an arousal response that may have affected the initial responses (Griskova et al., 2011; Griskova et al., 2007). Indeed, this period is often coincident with increased exploration of the recording chamber that dwindles over the following 20-30 min period (Drake et al., 1991).

We used female rats but did not monitor the estrous cycle during the study. While there was some session-to-session variability, it was not significant at the group level. It is however not possible to conclude that estrous cycling did not affect the measures in question. This is something that needs studying in future along with determining psychometric reliability in males.

There was no effect of session or session x time on the EEG measures. Since exploratory analysis suggested that response at time zero drove the time effect, we decided to omit sampling at time zero in the pharmacological study, to simplify data interpretation. This appeared to have removed time as a significant factor.

Our results indicate that while tone-evoked gamma as well as 40 Hz train-evoked gamma responses show acceptable reliability, 40 Hz ASSR is superior. Thus, at the group level, 40 Hz ASSR had a lower re-test related variance, compared to tone-evoked gamma. Since the same group of rats were repeatedly tested, we had an opportunity to evaluate if for a given subject, one measure evoked a more consistent response. We found that across ~ 30 test occasions conducted with the same 12 subjects, ASSR showed a consistently smaller CV than tone-evoked gamma, suggesting that, the ASSR measure is less variable and more precise. This is consistent with the general observation that narrow band steady state responses have better signal-noise ratios than transient, broadband oscillations (McFadden et al., 2014).

For testing the effect of the open channel blocker MK801 on gamma oscillations, we chose to include a range of doses that were reported to span an estimated occupancy of ~ 20% to 70% of the available NMDA receptor pool (Fernandes et al., 2015). MK801 showed robust suppression of both the evoked measures at 0.05 mg/kg and 0.15 mg/kg doses. However, 0.025 mg/kg of MK801 was effective only against 40 Hz ASSR. The lower variance in the ASSR measure may have helped reveal this low dose effect.

In summary, we report that the test-retest reliability of two frequently used response markers of sensory cortical circuit function that lend themselves for use as pharmacodynamic biomarkers. While both display acceptable psychometric properties for use in preclinical drug discovery and are sensitive to disruption of NMDA neurotransmission, particularly in the context of repeated measures, within subject experimental designs, 40 Hz ASSR has a quantitative edge over tone-evoked gamma.

## Abbreviations

ASSR: auditory steady state response
CV: coefficient of variation
EEG: electroencephalography
ICC: intraclass correlation coefficient
NMDA: N-methy1 D-aspartic acid
RMS: root mean square

## Acknowledgement

The authors gratefully acknowledge the help provided in the initial stages of this project by Dr. Ying Liu, Assistant Professor at the Department of Biostatistics and Epidemiology, East Tennessee State University, Johnson City, TN.

## Notes

**Conflict of interest**. None.

### Competing Interest Statement

The authors have declared no competing interest.

### Summary of Updates

The name spellings of the first author corrected from "Mohammad" to "Muhammad" Ummear Raza.

## References

Brenner, C. A., Krishnan, G. P., Vohs, J. L., Ahn, W. Y., Hetrick, W. P., Morzorati, S. L., & O’Donnell, B. F. (2009). Steady state responses: Electrophysiological assessment of sensory function in schizophrenia. In Schizophrenia Bulletin (Vol. 35, Issue 6, pp. 1065–1077). Oxford Academic. https://doi.org/10.1093/schbul/sbp091

Buzsáki, G., & Wang, X.-J. (2012). Mechanisms of Gamma Oscillations. Annual Review of Neuroscience, 35(1), 203–225. https://doi.org/10.1146/annurev-neuro-062111-150444

Cardin, J. A., Carlén, M., Meletis, K., Knoblich, U., Zhang, F., Deisseroth, K., Tsai, L. H., & Moore, C. I. (2009). Driving fast-spiking cells induces gamma rhythm and controls sensory responses. Nature, 459(7247), 663–667. https://doi.org/10.1038/nature08002

Cervenka, M. C., Franaszczuk, P. J., Crone, N. E., Hong, B., Caffo, B. S., Bhatt, P., Lenz, F. A., & Boatman-Reich, D. (2013). Reliability of early cortical auditory gamma-band responses. Clinical Neurophysiology, 124(124), 70–82. https://doi.org/10.1016/j.clinph.2012.06.003

Cleophas, T. J. M., & De Vogel, E. M. (1998). Crossover studies are a better format for comparing equivalent treatments than parallel-group studies. Pharmacy World and Science, 20(3), 113–117. https://doi.org/10.1023/A:1008626002664

Drake, M. E., Pakalnis, A., Phillips, B., Padamadan, H., & Hietter, S. A. (1991). Auditory Evoked Potentials in Anxiety Disorder. Clinical EEG and Neuroscience, 22(2), 97–101. https://doi.org/10.1177/155005949102200209

Fernandes, A., Wojcik, T., Baireddy, P., Pieschl, R., Newton, A., Tian, Y., Hong, Y., Bristow, L., & Li, Y. W. (2015). Inhibition of in vivo [3H]MK-801 binding by NMDA receptor open channel blockers and GluN2B antagonists in rats and mice. European Journal of Pharmacology, 766, 1–8. https://doi.org/10.1016/j.ejphar.2015.08.044

Gonzalez-Burgos, G., & Lewis, D. A. (2008). GABA Neurons and the Mechanisms of Network Oscillations: Implications for Understanding Cortical Dysfunction in Schizophrenia. Schizophrenia Bulletin, 34(34), 944–961. https://doi.org/10.1093/schbul/sbn070

Gonzalez-Burgos, Guillermo, & Lewis, D. A. (2012). NMDA receptor hypofunction, parvalbumin-positive neurons, and cortical gamma oscillations in schizophrenia. Schizophrenia Bulletin, 38(5), 950–957. https://doi.org/10.1093/schbul/sbs010

Griskova-Bulanova, I., Ruksenas, O., Dapsys, K., Maciulis, V., & Arnfred, S. M. H. (2011). Distraction task rather than focal attention modulates gamma activity associated with auditory steady-state responses (ASSRs). Clinical Neurophysiology, 122(8), 1541–1548. https://doi.org/10.1016/j.clinph.2011.02.005

Griskova, I., Morup, M., Parnas, J., Ruksenas, O., & Arnfred, S. M. (2007). The amplitude and phase precision of 40 Hz auditory steady-state response depend on the level of arousal. Experimental Brain Research, 183(1), 133–138. https://doi.org/10.1007/s00221-007-1111-0

Guo, Y., Logan, H. L., Glueck, D. H., & Muller, K. E. (2013). Selecting a sample size for studies with repeated measures. BMC Medical Research Methodology, 13(1), 100. https://doi.org/10.1186/1471-2288-13-100

Ip, C. T., Ganz, M., Ozenne, B., Sluth, L. B., Gram, M., Viardot, G., l’Hostis, P., Danjou, P., Knudsen, G. M., & Christensen, S. R. (2018). Pre-intervention test-retest reliability of EEG and ERP over four recording intervals. International Journal of Psychophysiology, 134, 30–43. https://doi.org/10.1016/j.ijpsycho.2018.09.007

Javitt, D. C. (2009a). When doors of perception close: bottom-up models of disrupted cognition in schizophrenia. Annual Review of Clinical Psychology, 5, 249–275. https://doi.org/10.1146/annurev.clinpsy.032408.153502

Javitt, D. C. (2009b). Sensory processing in schizophrenia: Neither simple nor intact. In Schizophrenia Bulletin (Vol. 35, Issue 6, pp. 1059–1064). Oxford Academic. https://doi.org/10.1093/schbul/sbp110

Javitt, D. C., & Sweet, R. A. (2015). Auditory dysfunction in schizophrenia: Integrating clinical and basic features. In Nature Reviews Neuroscience (Vol. 16, Issue 9, pp. 535–550). Nature Publishing Group. https://doi.org/10.1038/nrn4002

Koo, T. K., & Li, M. Y. (2016). A Guideline of Selecting and Reporting Intraclass Correlation Coefficients for Reliability Research. Journal of Chiropractic Medicine, 15(2), 155. https://doi.org/10.1016/J.JCM.2016.02.012

Kozono, N., Honda, S., Tada, M., Kirihara, K., Zhao, Z., Jinde, S., Uka, T., Yamada, H., Matsumoto, M., Kasai, K., & Mihara, T. (2019). Auditory Steady State Response; nature and utility as a translational science tool. Scientific Reports, 9(1), 8454. https://doi.org/10.1038/s41598-019-44936-3

Krishnan, R. R., Kraus, M. S., & Keefe, R. S. E. (2011). Comprehensive model of how reality distortion and symptoms occur in schizophrenia: Could impairment in learning-dependent predictive perception account for the manifestations of schizophrenia? Psychiatry and Clinical Neurosciences, 65(65), 305–317. https://doi.org/10.1111/j.1440-1819.2011.02203.x

Lauschke, V. M., Zhou, Y., & Ingelman-Sundberg, M. (2019). Novel genetic and epigenetic factors of importance for inter-individual differences in drug disposition, response and toxicity. In Pharmacology and Therapeutics (Vol. 197, pp. 122–152). Elsevier Inc. https://doi.org/10.1016/j.pharmthera.2019.01.002

Legget, K. T., Hild, A. K., Steinmetz, S. E., Simon, S. T., & Rojas, D. C. (2017). MEG and EEG demonstrate similar test-retest reliability of the 40 Hz auditory steady-state response. International Journal of Psychophysiology, 114, 16–23. https://doi.org/10.1016/j.ijpsycho.2017.01.013

Leicht, G., Karch, S., Karamatskos, E., Giegling, I., Möller, H. J., Hegerl, U., Pogarell, O., Rujescu, D., & Mulert, C. (2011). Alterations of the early auditory evoked gamma-band response in first-degree relatives of patients with schizophrenia: Hints to a new intermediate phenotype. Journal of Psychiatric Research, 45(5), 699–705. https://doi.org/10.1016/j.jpsychires.2010.10.002

Leicht, G., Vauth, S., Polomac, N., Andreou, C., Rauh, J., Mußmann, M., Karow, A., & Mulert, C. (2016). EEG-Informed fMRI Reveals a Disturbed Gamma-Band-Specific Network in Subjects at High Risk for Psychosis. Schizophrenia Bulletin, 42(1), 239–249. https://doi.org/10.1093/schbul/sbv092

Light, G. A., Joshi, Y. B., Molina, J. L., Bhakta, S. G., Nungaray, J. A., Cardoso, L., Kotz, J. E., Thomas, M. L., & Swerdlow, N. R. (2020). Neurophysiological biomarkers for schizophrenia therapeutics. Biomarkers in Neuropsychiatry, 2, 100012. https://doi.org/10.1016/j.bionps.2020.100012

Luck, S. J., Mathalon, D. H., O’Donnell, B. F., Hmlinen, M. S., Spencer, K. M., Javitt, D. C., & Uhlhaas, P. J. (2011). A roadmap for the development and validation of event-related potential biomarkers in schizophrenia research. In Biological Psychiatry (Vol. 70, Issue 1, pp. 28–34). Elsevier USA. https://doi.org/10.1016/j.biopsych.2010.09.021

McFadden, K. L., Steinmetz, S. E., Carroll, A. M., Simon, S. T., Wallace, A., & Rojas, D. C. (2014). Test-Retest Reliability of the 40 Hz EEG Auditory Steady-State Response. PLoS ONE, 9(1), e85748. https://doi.org/10.1371/journal.pone.0085748

Molina, J. L., Thomas, M. L., Joshi, Y. B., Hochberger, W. C., Koshiyama, D., Nungaray, J. A., Cardoso, L., Sprock, J., Braff, D. L., Swerdlow, N. R., & Light, G. A. (2020). Gamma oscillations predict pro-cognitive and clinical response to auditory-based cognitive training in schizophrenia. Translational Psychiatry, 10(1), 1–10. https://doi.org/10.1038/s41398-020-01089-6

Oribe, N., Hirano, Y., del Re, E., Seidman, L. J., Mesholam-Gately, R. I., Woodberry, K. A., Wojcik, J. D., Ueno, T., Kanba, S., Onitsuka, T., Shenton, M. E., Goldstein, J. M., Niznikiewicz, M. A., McCarley, R. W., & Spencer, K. M. (2019). Progressive reduction of auditory evoked gamma in first episode schizophrenia but not clinical high risk individuals. Schizophrenia Research, 208, 145–152. https://doi.org/10.1016/j.schres.2019.03.025

Perez, V. B., Roach, B. J., Woods, S. W., Srihari, V. H., McGlashan, T. H., Ford, J. M., & Mathalon, D. H. (2013). Early auditory gamma-band responses in patients at clinical high risk for schizophrenia. In Supplements to Clinical Neurophysiology (Vol. 62, pp. 147–162). Elsevier B.V. https://doi.org/10.1016/B978-0-7020-5307-8.00010-7

Rissling, A. J., & Light, G. A. (2010). Neurophysiological measures of sensory registration, stimulus discrimination, and selection in schizophrenia patients. In Current Topics in Behavioral Neurosciences (Vol. 4, pp. 283–309). Springer Verlag. https://doi.org/10.1007/7854_2010_59

Roach, B. J., D’Souza, D. C., Ford, J. M., & Mathalon, D. H. (2019). Test-retest reliability of time-frequency measures of auditory steady-state responses in patients with schizophrenia and healthy controls. NeuroImage: Clinical, 23, 101878. https://doi.org/10.1016/j.nicl.2019.101878

Roach, B. J., Ford, J. M., & Mathalon, D. H. (2019). Gamma Band Phase Delay in Schizophrenia. Biological Psychiatry: Cognitive Neuroscience and Neuroimaging, 4(2), 131–139. https://doi.org/10.1016/j.bpsc.2018.08.011

Ross, B., Schneider, B., Snyder, J. S., & Alain, C. (2010). Biological markers of auditory gap detection in young, middle-aged, and older adults. PLoS ONE, 5(4), 10101. https://doi.org/10.1371/journal.pone.0010101

Salkind, N. (2010). Encyclopedia of Research Design. In Encyclopedia of Research Design. SAGE Publications, Inc. https://doi.org/10.4135/9781412961288

Silverstein, S. M., & Keane, B. P. (2011). Vision science and schizophrenia research: Toward a re-view of the disorder editors’ introduction to special section. In Schizophrenia Bulletin (Vol. 37, Issue 4, pp. 681–689). Schizophr Bull. https://doi.org/10.1093/schbul/sbr053

Sivarao, D. V. (2015). The 40-Hz auditory steady-state response: A selective biomarker for cortical NMDA function. Annals of the New York Academy of Sciences, 1344(1344), 27–36. https://doi.org/10.1111/nyas.12739

Sivarao, D. V, Chen, P., Senapati, A., Yang, Y., Fernandes, A., Benitex, Y., Whiterock, V., Li, Y.-W., & Ahlijanian, M. K. (2016). 40 Hz Auditory Steady-State Response Is a Pharmacodynamic Biomarker for Cortical NMDA Receptors. Neuropsychopharmacology, 41(9), 2232. https://doi.org/10.1038/NPP.2016.17

Spellman, T. J., & Gordon, J. A. (2015). Synchrony in schizophrenia: A window into circuit-level pathophysiology. In Current Opinion in Neurobiology (Vol. 30, pp. 17–23). Elsevier Ltd. https://doi.org/10.1016/j.conb.2014.08.009

Stroup, T. S., Lieberman, J. A., McEvoy, J. P., Swartz, M. S., Davis, S. M., Capuano, G. A., Rosenheck, R. A., Keefe, R. S. E., Miller, A. L., Belz, I., & Hsiao, J. K. (2007). Effectiveness of olanzapine, quetiapine, and risperidone in patients with chronic schizophrenia after discontinuing perphenazine: A CATIE study. American Journal of Psychiatry, 164(164), 415–427. https://doi.org/10.1176/ajp.2007.164.3.415

Tan, H. R. M., Gross, J., & Uhlhaas, P. J. (2015). MEG-measured auditory steady-state oscillations show high test-retest reliability: A sensor and source-space analysis. NeuroImage, 122, 417–426. https://doi.org/10.1016/j.neuroimage.2015.07.055

Tiesinga, P., & Sejnowski, T. J. (2009). Cortical Enlightenment: Are Attentional Gamma Oscillations Driven by ING or PING? In Neuron (Vol. 63, Issue 6, pp. 727–732). Neuron. https://doi.org/10.1016/j.neuron.2009.09.009

